# When can site-specific glycoform compositions resolve biosynthetic rates?

**DOI:** 10.64898/2026.10.09.757869

**Authors:** Yusuke Matsui, Koichi Himori, Bingyuan Zhang, Akihiro Kitani

## Abstract

Mass spectrometry measures site-specific glycoform compositions, but when these measurements can resolve upstream biosynthetic rates remains unclear. We assess whether small changes in effective rates, each representing the processing capacity of a group of biosynthetic reactions, can be distinguished near specified baseline rates with secretion held fixed. Because proportions sum to one, *m* jointly normalised glycoform classes provide at most *m* − 1 independent quantities. Distinguishing changes in *k* rates therefore requires at least *k* + 1 classes, although this count alone does not ensure identifiability. In a canonical N-glycan network with eight rates, exhaustive evaluation of nine-class panels identified panels that could locally distinguish all eight rates. Among these panels, class choice changed the allowable measurement error by approximately 18-fold for the same target uncertainty in inferred rates. A separate simulation within the fixed network showed that estimating rate-change combinations with weaker composition responses reduced bias but amplified measurement noise. We applied the framework to plasma data from 40 controls and 70 patients across 14 inherited glycosylation-disorder groups. A prespecified composition score for the expected pathway disturbance ranked among the top two in 13 of 14 groups. Network responses linked observed composition shifts to biosynthetic reaction classes. Expanding site models from five to eight reaction classes improved control-composition fit but reduced the number of sites at which all modelled rates were locally distinguishable and increased noise sensitivity. Rate inference thus requires sufficient panel size, separable rate responses, adequate measurement precision and a model that fits the composition. These requirements guide measurement design and help determine whether compositions support pathway-level interpretation or individual-rate inference.

**Author summary:** Mass spectrometry measures the relative amounts of glycan forms attached to a protein site. We asked when these proportions can distinguish changes in the biosynthetic reactions that produce them. Because proportions sum to 100%, a panel contains one fewer independent quantity than its number of glycoform classes. Our eight-rate model therefore required at least nine classes. Which classes were measured strongly affected the measurement precision needed to distinguish the rates. We applied the framework to plasma data from inherited glycosylation disorders. The expected composition change ranked among the two strongest signals in 13 of 14 disorder groups. Network responses linked these changes to reaction groups. Most site models could distinguish small rate changes near a fitted control reference, but this did not establish recovery of patients’ true rates. A more detailed model reproduced compositions better while making individual rates harder to distinguish. The framework helps researchers select informative glycoforms, calculate measurement-precision requirements and assess which biosynthetic changes the data can support. Its conclusions depend on the assumed network, reference rates and measurement-error model.

## Introduction

Protein glycosylation generates molecular forms of the same protein that carry different glycans at the same attachment site [1, 2]. Their relative abundances constitute a site-specific glycoform composition. This composition is an output of glycan processing in the endoplasmic reticulum and Golgi apparatus and depends on how rapidly each reaction proceeds relative to other reactions and to protein trafficking and secretion. Processing is influenced not only by enzyme activity but also by donor supply and transport and site accessibility [1, 3]. Understanding a composition change therefore requires examining which changes in biosynthetic processing could explain it. Mass spectrometry can quantify large numbers of site-specific glycoforms [4–6], but the plasma data considered here report relative within-site compositions, not the intracellular reaction rates themselves (Fig 1A).

**Fig 1.**
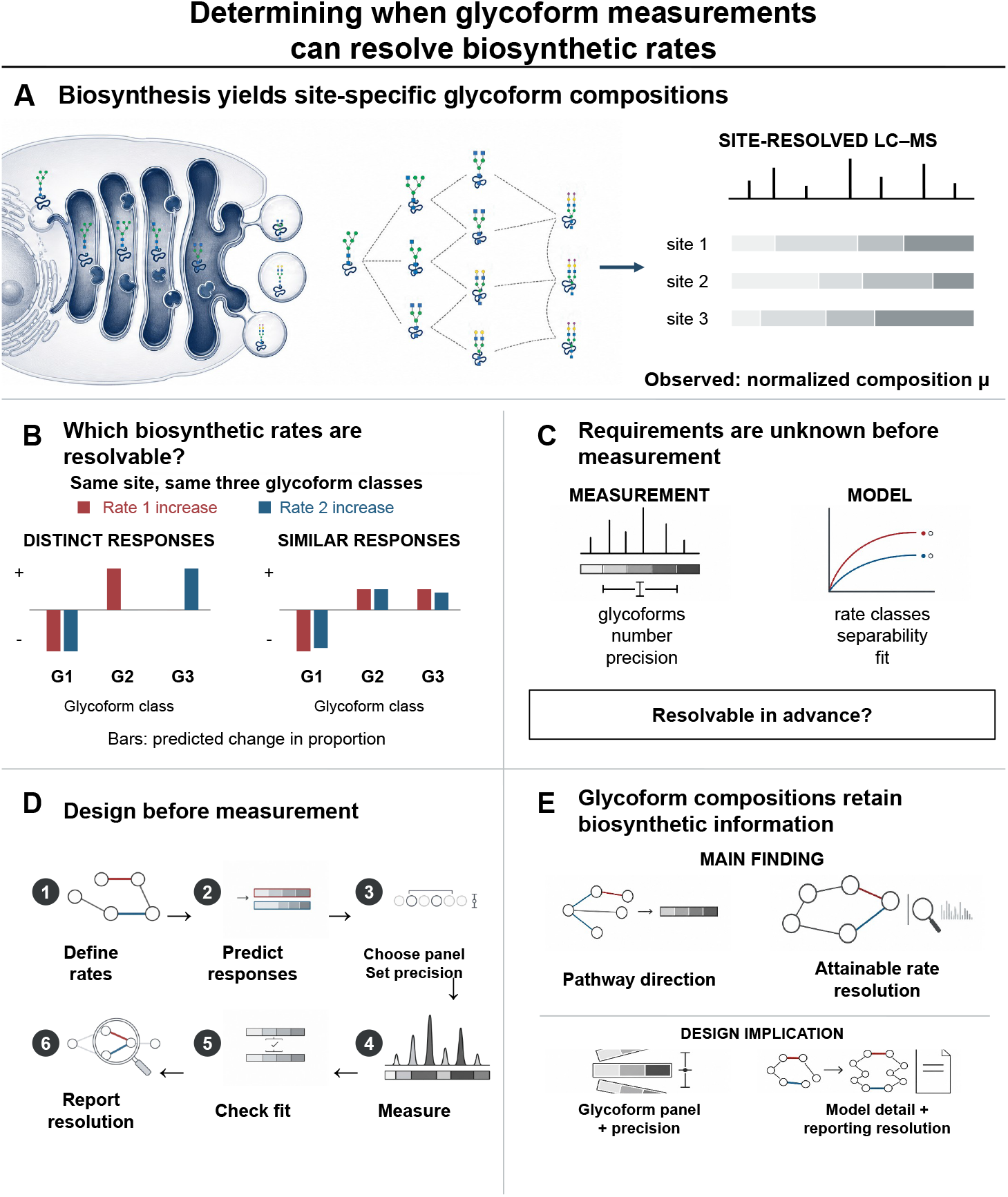
Determining when glycoform measurements can resolve biosynthetic rates. (A) Liquid chromatography–mass spectrometry (LC–MS) measures the normalised glycoform composition at each site. (B) Red and blue bars show schematic changes in the same three glycoform proportions after a small increase in either of two effective rates. G1–G3 are generic class labels, not structural assignments; each response sums to zero. Linearly independent responses distinguish the rates locally, whereas nearly proportional responses are difficult to separate under noise. (C) Feasibility depends on glycoform number, measurement precision, rate-class separability and model fit. (D) Define target rates, predict responses, choose a panel and required precision, measure, check fit and report the supported biosynthetic detail. (E) Compositions retain pathway information and may resolve individual rate directions; measurement design and model detail should match the intended inference. Glycan drawings follow the Symbol Nomenclature for Glycans (SNFG). Spectra, networks and response bars illustrate the study logic rather than quantitative results. Non-data conceptual artwork and icons were prepared with OpenAI image generation; scientific labels and response diagrams were checked separately.

Reaction-network models and pathway-mapping studies link biosynthetic processing to measured glycan profiles [7–10]. Models have also been used to estimate enzyme activities or concentrations from those profiles [11, 12], and recent work has assessed whether the available data constrain those estimates [13]. We build on these studies by connecting measurement design to the biosynthetic information that can be inferred: which glycoform classes must be measured, how precisely, and how much detail about reaction-rate changes those measurements can support.

For example, in a sequential pathway, can the selected glycoform proportions distinguish slower upstream processing from slower downstream processing (Fig 1B)?

Here, an *effective rate* represents the aggregate processing capacity of a group of reactions and may reflect enzyme activity, donor availability, transport and other pathway influences [1, 3, 14, 15]. A change in an effective rate does not uniquely identify the responsible enzyme or gene.

Distinguishing changes in effective rates requires differences in the composition responses produced by those changes. These differences must first be present in the selected measurements. Distinct glycan states may be combined into the same measured class, and proportions normalised across the measured classes sum to one, limiting the number of independent quantities. If a rate change leaves the measured proportions unchanged, or different combinations of rate changes produce the same response, increasing measurement precision cannot distinguish them. Even when responses differ, small differences can make rate estimates unstable under measurement error. We therefore assess two questions separately: can rate changes be resolved without measurement error, and how much uncertainty does measurement error introduce? A model that reproduces the observed composition well does not necessarily meet either requirement (Fig 1C).

These assessments inform measurement design and biological interpretation. Before data collection, responses predicted under an assumed network and baseline rates can guide the choice of measurements and precision needed to distinguish target reactions. After data collection, we check composition fit and assess whether the data support pathway-level interpretation or individual effective-rate inference (Fig 1D). We compared measurement panels and ran simulations in a canonical N-glycan network, then applied the framework to public plasma glycoproteomic data from inherited glycosylation disorders [16]. Using disorders with known defects in glycan processing, donor synthesis or transport [17, 18], we tested whether composition scores captured expected pathway changes, whether network responses explained disease-associated composition shifts, and whether individual rate changes were locally distinguishable. We also compared coarser and more detailed networks to assess the relationship between composition fit and rate identifiability (Fig 1E). We evaluate how well small changes from baseline rates can be inferred under a specified network and measurement-error model.

## Model and definitions

Assessing what a measured composition can reveal about biosynthesis requires a model that connects reaction rates to observed glycoform classes. We define that connection here, together with the local response used to distinguish rate changes.

### Glycan states and reaction rules

At one glycosylation site, an *observation class* groups glycoforms by measured glycan composition. Hex_5_HexNAc_2_ (H5N2 in Fig. 2) denotes five hexose and two N-acetylhexosamine residues. Composition labels do not specify linkage or branching structures. A network node represents a modelled glycan state; several states can share one observed composition. Directed edges represent allowed processing reactions, with substrate conditions and reaction order based on N-glycan biosynthesis [19] and earlier mechanistic and rule-based models [7, 8, 11].

**Fig 2.**
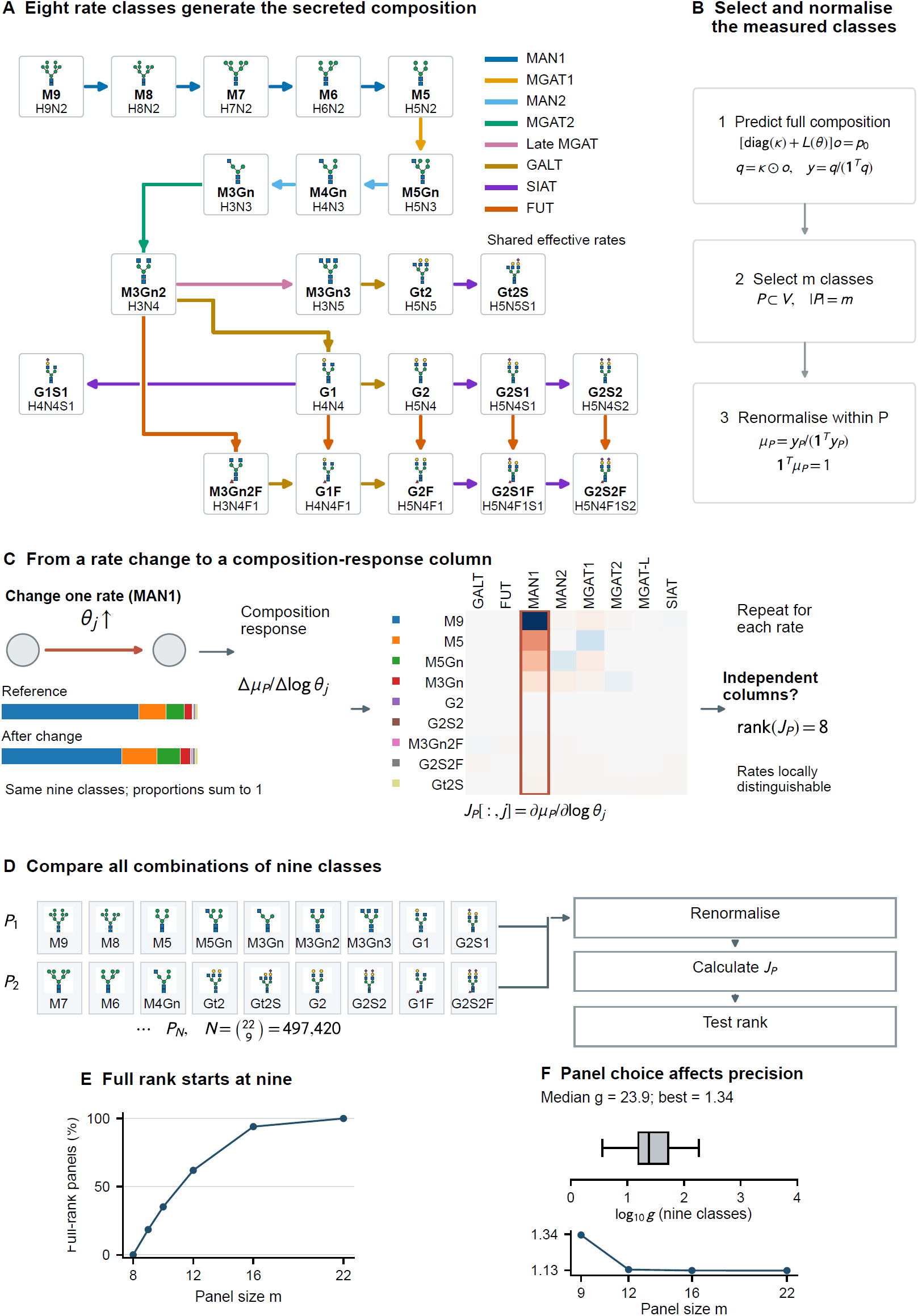
Nine classes are necessary, but panel choice determines rate recoverability in the canonical graph. (A) The 22-state N-glycan graph has eight shared effective rates (coloured edges), Man9 entry and fixed secretion. Late MGAT is abbreviated MGAT-L in C. (B) Selecting and renormalising *m* classes leaves at most *m* − 1 independent composition directions. (C) Changing one rate alters the panel composition, generating one column of *J*_*P*_ . The bars illustrate a 0.2 increase in log MAN1 rate for the same nine classes; the heatmap shows the local derivatives at unit rates for the best nine-class panel (red, positive; blue, negative), with the MAN1 column outlined. Rank 8 requires eight independent response columns. (D) Two example nine-class panels illustrate the enumeration. Each panel is renormalised before calculating *J*_*P*_ and testing its rank. All 497,420 combinations were tested. (E) No panel with ≤ 8 attained rank 8; 92,191 nine-class panels (18.5%) did at tolerance 10^*−*9^.Points show enumerated sizes; lines guide the eye. (F) For full-rank nine-class panels, median *g* = 23.9 and best *g* = 1.34. Boxes, central lines and whiskers show the 25th–75th percentiles, median and 5th–95th percentiles of log_10_ *g*. The best *g* approaches 1.13 at larger panel sizes. Smaller *g* means less noise amplification. Glycan drawings follow SNFG; H, N, F and S denote Hex, HexNAc, Fuc and Neu5Ac. All results are local and conditional on the specified model.

For example, mannose trimming connects Hex_6_HexNAc_2_ to Hex_5_HexNAc_2_, whereas galactose addition to a suitable acceptor connects Hex_3_HexNAc_4_ to Hex_4_HexNAc_4_ (Fig 2A). A matching composition difference alone does not create an edge: the modelled substrate must also satisfy the reaction rule. For instance, in the site models, galactose addition requires an unfilled antenna and sialic-acid addition requires a galactose not yet sialylated. These coarse rules constrain possible processing without assigning a unique structure to each measured composition. Reaction rules and state-generation limits are detailed in Methods, “Network construction”, and S1 Text, Supplementary Methods S3.

A *rate class* groups edges sharing one positive effective rate, such as the galactose-addition reactions in the GALT class. The vector *θ* = (*θ*_1_, …, *θ*_*k*_) contains the *k* rates present in a network. These are relative processing rates, not measured enzyme abundances or absolute biochemical rate constants.

### From reaction rates to secreted composition

Let *W*_*ij*_(*θ*) be the rate of the directed reaction from state *i* to state *j*, and let *c*(*i* → *j*) identify its rate class. The reaction matrix is constructed as

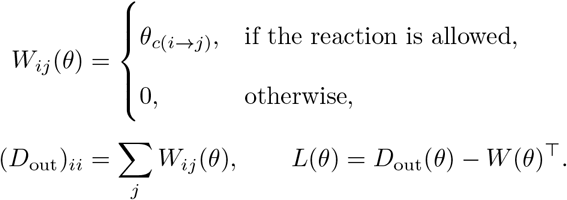

Here, *D*_out_ is diagonal: it accounts for total reaction outflow from each state, while *W*^*⊤*^ accounts for inflow from other states. Processing and secretion are modelled as first-order transitions in a single well-mixed pool. Glycans enter through a fixed input vector *p*_0_, undergo processing and leave each state at a fixed positive secretion rate *κ*_*i*_. At steady state, external input and reaction inflow balance reaction outflow and secretion:

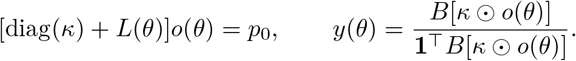

The vector *o*(*θ*) gives the steady-state amount of each glycan state. Changing the processing rates *θ* changes the reaction matrix *L*(*θ*), and hence these amounts and the secretion fluxes *κ* ⊙ *o*(*θ*); ⊙ denotes elementwise multiplication. The observation matrix *B* maps modelled states to measured classes: *B*_*ci*_ = 1 if state *i* belongs to class *c*, and is zero otherwise. For example, two states with different glycan structures but the same measured composition contribute to the same class, so their secretion fluxes are added. Dividing these class fluxes by their total gives the predicted composition *y*(*θ*), with **1** denoting a vector of ones. In the canonical analysis, each state is treated as a separate observation class and *B* is the identity matrix. Fixing secretion sets the relative timescale: multiplying all processing and secretion rates by the same factor leaves the composition unchanged.

For each rate vector, we solve the linear steady-state equations, aggregate the secretion fluxes and normalise them. Small rate perturbations then give numerical composition sensitivities. The solvers and differentiation steps are specified in Methods, “Forward calculation and numerical sensitivities”; the steady-state proof and empirical sensitivity calculation are in S1 Text, Supplementary Methods S3 and S4, respectively.

### Observation panels and local rate identifiability

An *observation panel* contains classes normalised together. In a hypothetical Hex_5_HexNAc_2_, Hex_6_HexNAc_2_, Hex_7_HexNAc_2_ panel with proportions of 20%, 30% and 50%, two proportions determine the third, leaving two independently varying quantities. These illustrative percentages are not study measurements. For a panel *P* of *m* classes with positive total predicted abundance, class *i* has predicted proportion

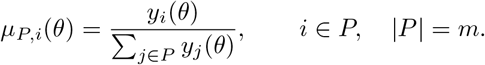

We evaluate composition responses near a baseline rate vector *θ*_0_, termed the operating point. These baseline rates are specified in the canonical panel analysis and fitted to the mean control composition in each empirical site model. We express changes on the log-rate scale so that sensitivity describes relative changes in processing rates rather than absolute increments. For a small change, Δ log *θ*_*j*_ ≈ Δ*θ*_*j*_*/θ*_0,*j*_; thus, a 10% increase represents the same log-rate change regardless of the baseline rate. The local composition response is

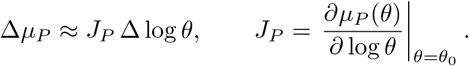

Each column of the *m* × *k* Jacobian *J*_*P*_ describes the composition response to a small relative change in one rate. A rate direction is a pattern of changes in one or more log rates. If a nonzero combination of rate changes produces no first-order composition change, its effects cancel and cannot be distinguished from no change by this local approximation. More generally, adding that combination to another rate change gives the same predicted response. Full column rank excludes such cancellation: the columns are linearly independent, allowing all *k* log-rate changes to be distinguished within the local linear approximation.

Because the proportions sum to one, each response column sums to zero.

Consequently,

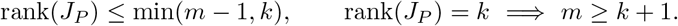

The class-count requirement is necessary but does not guarantee full rank. Even at full rank, weak or nearly overlapping responses can make rate estimates highly sensitive to measurement noise. We use singular-value decomposition to assess the number and strength of independent response directions, and propagate composition error through the local inverse to evaluate rate uncertainty. In the empirical analyses, each panel represents one peptide-defined glycosylation site. Fitting is detailed in Methods, “Control-composition fitting and model comparison” (S1 Text, Supplementary Methods S4). Rank thresholds and error propagation are specified under “Local rank and conditioning” and “Measurement-error amplification”, respectively (S1 Text, Supplementary Methods S4; Tables S8–S13).

We use this framework to address three questions.

1. **Which observation panels can resolve local rate changes, and at what measurement precision?** We examine how the number and choice of glycoform classes determine independent rate responses and the amplification of measurement error (Fig 2).
2. **What biosynthetic information is retained in measured plasma compositions?** We test whether composition scores capture expected pathway disturbances and whether network responses account for disease-associated shifts, then assess local rate identifiability in the fitted site models (Figs 3–4).
3. **How does the detail retained in rate estimation affect fit and uncertainty?** We distinguish two analyses: retaining additional sensitivity modes within a fixed network, and expanding the number of biochemical reaction classes in the site models (Fig 5).

**Fig 3.**
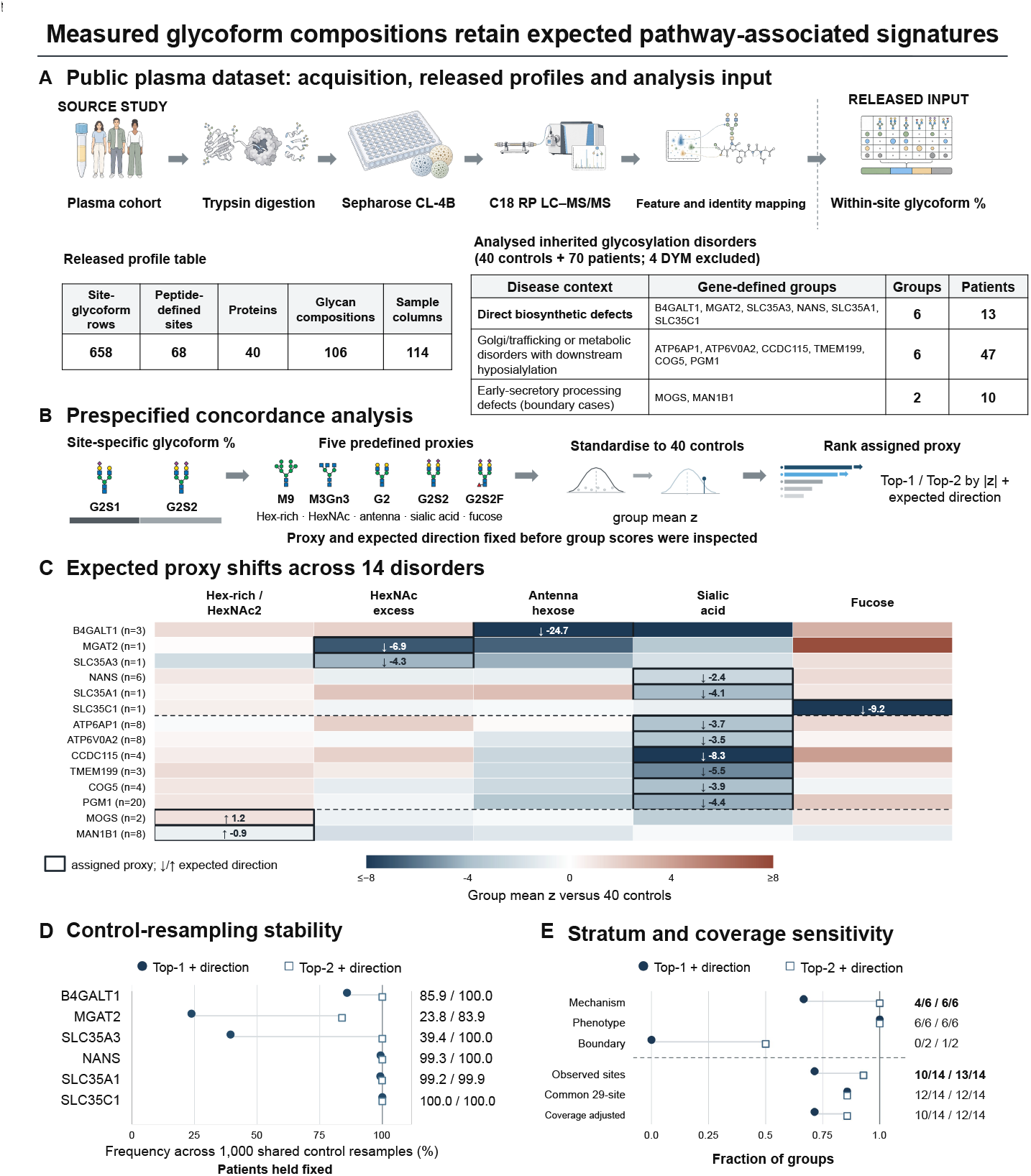
Measured glycoform compositions retain expected pathway-associated signatures. (A) Public PXD034214 acquisition workflow, with non-data workflow illustrations prepared using OpenAI image generation, released profile dimensions and analysed cohort: 40 controls and 70 patients in 14 inherited glycosylation-disorder groups after exclusion of four DYM-labelled samples. (B) Prespecified concordance workflow. Site-specific percentages were summarised by five fixed pathway scores, standardised against the controls and ranked by the absolute group-mean *z* score; sign concordance was evaluated separately. Assignments and directions were fixed before group scores were inspected. Glycan drawings illustrate structures consistent with the measured compositions, which do not uniquely resolve linkage, branching isomers or fucose position. (C) Group-mean *z* scores across the 14 disorders. Boxes mark the target score, arrows show the expected direction, colours are clipped at ±8, and dashed lines separate the three strata. (D) Control-resampling frequency of Top-1 and Top-2 expected-direction concordance in the six mechanism-anchored disorders. Patients were held fixed, so this measures control-calibration stability rather than patient-population uncertainty. (E) Concordance by stratum and after restriction to 29 common sites or control-based coverage adjustment. The six phenotype-anchored groups share one downstream sialylation proxy and are not independent causal-mechanism recoveries.

**Fig 4.**
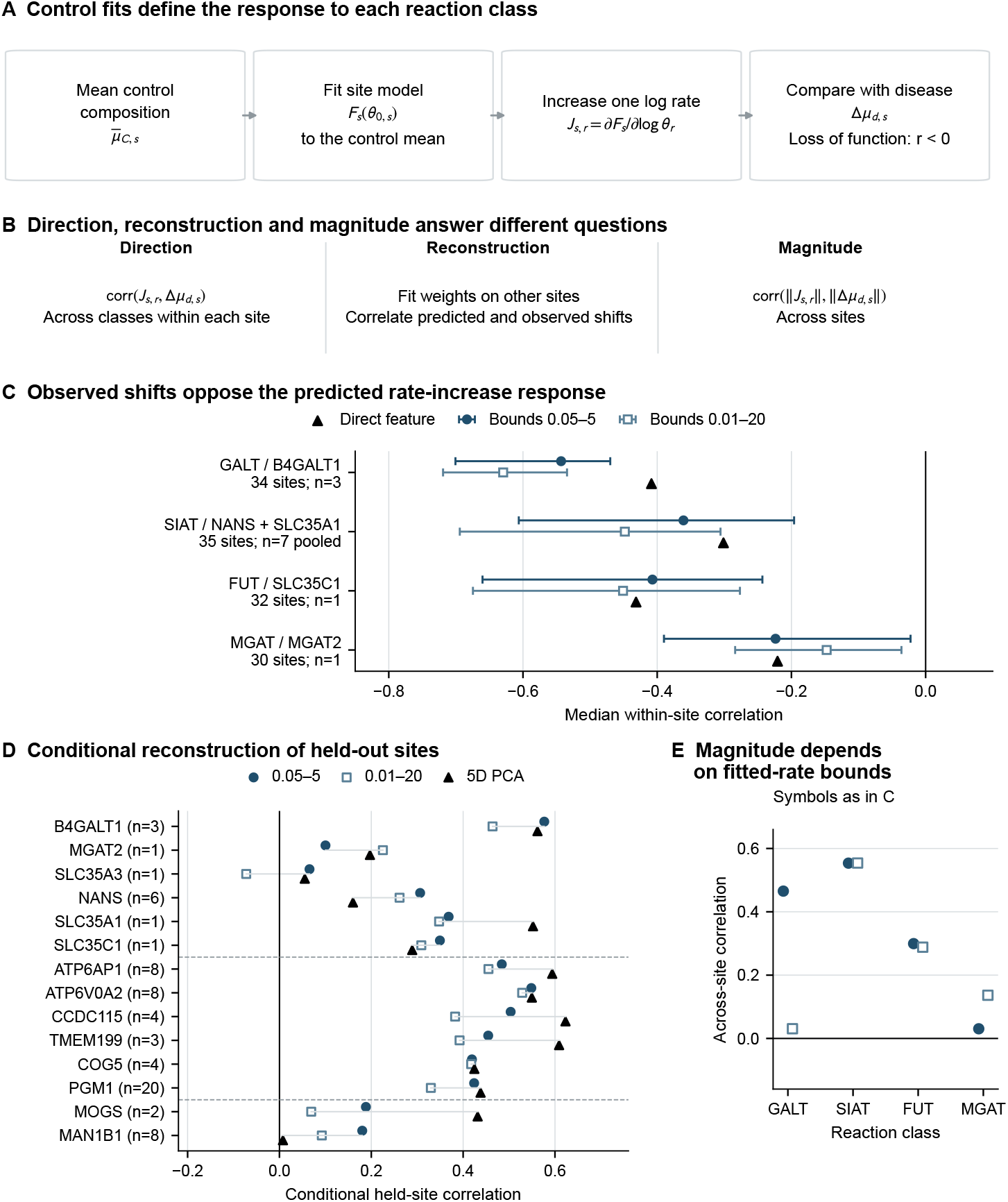
Network sensitivities align site-specific glycoform shifts with expected loss-of-function directions. (A) The mean control composition at each site was fitted to define its operating point. The log-rate sensitivity *J*_*r*_ predicts the glycoform shift caused by increasing reaction class *r* and was compared with the observed loss-of-function shift Δ*µ*; negative correlation is therefore expected. (B) Separate analyses test within-site direction, conditional reconstruction of a held-site response and across-site magnitude. (C) Median direction correlations for a support-matched direct feature and for the same network fitted under rate bounds of 0.05–5 or 0.01–20. Error bars are pointwise 95% protein-cluster resampling intervals, not patient-population intervals. SIAT and FUT represent lumped sialylation and fucosylation capacities. (D) Disorder-specific conditional leave-one-site-out correlations for the two network fits and a five-component principal-component basis. This is not patient-level cross-validation.(E) Across-site magnitude correlations under the two rate bounds. Both fitting bounds apply to the same five-class model; this is separate from the eight-class augmentation. The wider interval tests fitting sensitivity, not a biological range. Network analyses use graph-supported conditional compositions; unsupported mass is reported separately.

**Fig 5.**
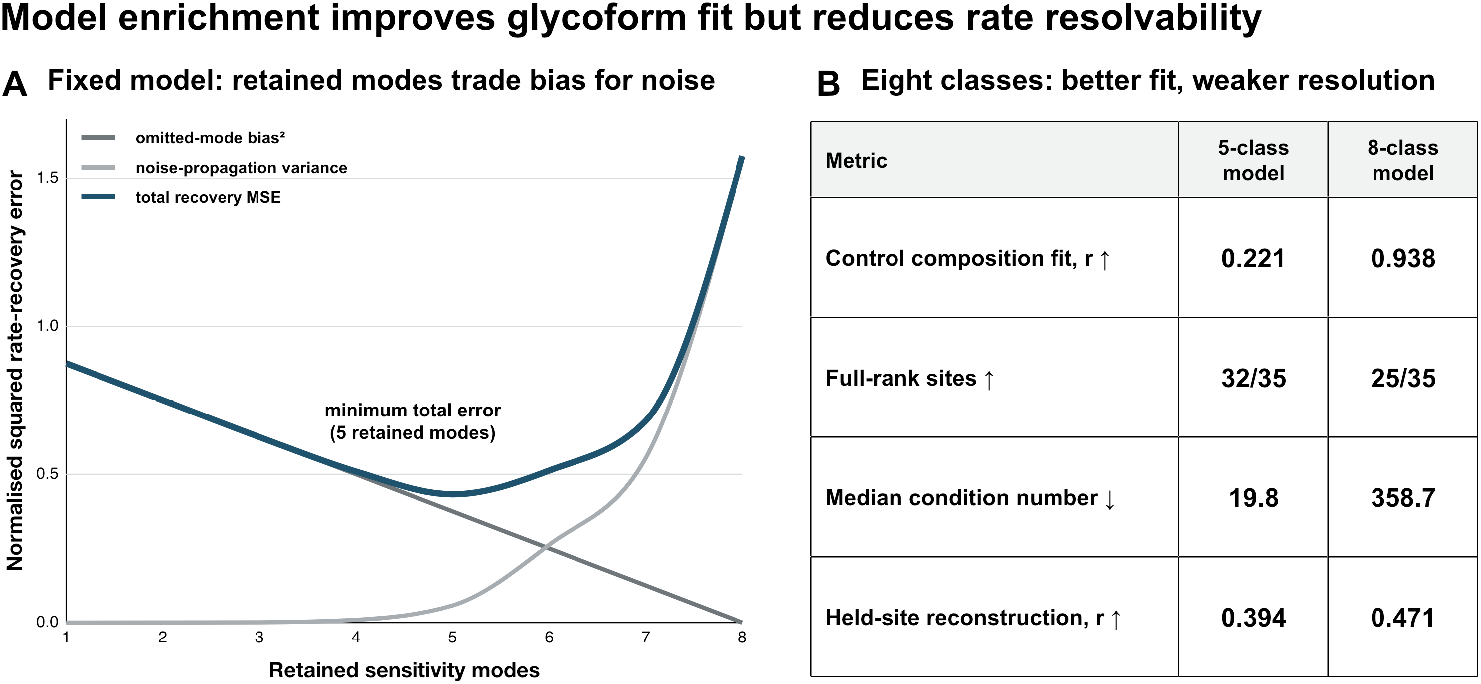
Model enrichment improves glycoform fit but reduces rate resolvability. (A) In a simulation within the fixed canonical eight-class model, retaining additional sensitivity modes reduced bias from omitted log-rate directions but increased variance propagated from measurement noise. Their sum, the normalised mean-squared rate-recovery error, was lowest at five retained modes. Curves show medians across 2,048 operating points at an assumed multinomial total count of 300,000, which sets the simulated composition noise. Retained modes describe estimator complexity and are not the empirical five reaction classes. (B) In empirical refits of 35 sites, expanding from five to eight reaction classes markedly improved control-composition fit and modestly improved conditional held-site reconstruction, but reduced the number of full-rank sites and increased the median condition number. Fitted rates were constrained to 0.05–5. Held-site reconstruction was evaluated within disorder and does not represent patient-level prediction.

## Results

### The canonical eight-rate model requires at least nine glycoform classes per panel, and panel choice controls recoverability

The class-count requirement alone does not determine which combinations of glycoforms can distinguish rates. We therefore used a specified canonical network to evaluate how panel selection affects local identifiability and precision of rate recovery.

The canonical network comprised 22 states and 25 directed reactions, with Man9 entry and fixed uniform secretion (Fig 2A). Its eight rate classes distinguish two mannose-trimming stages, three HexNAc-addition stages, galactosylation, sialylation and fucosylation. This was a model-based panel-design calculation. We set all eight processing rates to 1 and the secretion rate at every state to 0.5, using the same reference setting for every panel; these rates were specified rather than fitted to patient or control data.

For each candidate panel, we selected classes from the predicted steady-state output and renormalised their proportions to sum to one. A panel of *m* classes therefore contains at most *m* − 1 independently varying quantities, so distinguishing eight rate directions requires at least nine classes. We then perturbed each log rate separately around the reference setting and calculated the changes in the panel proportions. The eight response patterns form the columns of the panel Jacobian *J*_*P*_, defined in “Model and definitions”. Singular-value decomposition tested whether these columns were linearly independent, giving rank 8 (Fig 2A–C). We tested every combination of nine classes selected from the graph’s 22 states, giving 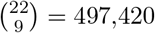 candidate panels. Of these, 92,191 (18.5%) attained rank 8 at the evaluated operating point and could therefore distinguish all eight rate directions locally (Fig 2D,E). This count used an absolute singular-value tolerance of 10^*−*9^; changing the tolerance changes the qualifying count (Methods, “Local rank and conditioning”; Table S3). The remaining panels were numerically rank deficient at that tolerance. Stability under measurement error required a separate assessment.

To assess precision, we assumed independent relative errors with the same measurement coefficient of variation (CV) for each class before panel normalisation. We propagated these errors through normalisation and the local inverse of the composition response. The noise-amplification factor *g* is the standard deviation per unit CV in the most uncertain unit-length direction in log-rate space. Thus, for a target standard deviation *δ*,

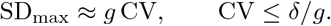

Smaller *g* means less error amplification; the best panel is the one with the smallest *g* at the specified reference rates. The covariance calculation is detailed in Methods, “Measurement-error amplification”.

Among the qualifying nine-class panels, *g* had a median of 23.9 and a best value of 1.34 (Fig 2F). For an illustrative, analyst-selected target standard deviation of inferred log rates of *δ* = 0.25, the median panel required an analytical coefficient of variation (CV) no greater than 1.05%, whereas the best panel tolerated 18.7%, a 17.9-fold difference. The target is not a biologically validated error limit; alternative targets of 0.10, 0.50 and 1.00 are compared in Table S11. Within the local error model, allowable CV scales linearly with the chosen target, so the ratio between panels does not depend on choosing 0.25. Here, CV is the measurement-error standard deviation divided by the mean abundance; smaller CV means higher precision. Increasing panel size provided much less benefit: the best *g* improved only from 1.34 at nine classes to 1.13 at 12 classes and changed little at 16 and 22 classes (Fig S4A,B; Tables S11 and S12).

For this graph, rate recovery requires at least nine glycoform classes chosen to give independent responses with limited noise amplification. These CV thresholds are design calculations under the assumed measurement-error model, not measured assay performance. The requirements are local and conditional on the graph and operating point; rank and precision calculations are described in Methods, “Local rank and conditioning” and “Measurement-error amplification”. Canonical rank checks are in S1 Text, Supplementary Results S6 (Fig S1A; Table S3), and precision calculations are documented in Tables S11–S13.

### Plasma glycoform compositions retain expected pathway-associated signatures of inherited glycosylation defects

Before assessing rate inference in the plasma data, we asked whether the measured compositions retained changes associated with known biosynthetic defects. We used PXD034214 as a retrospective positive-control dataset to compare prespecified composition scores with the expected pathway changes.

The author-released, preprocessed dataset contained within-site glycoform percentages at 68 peptide-defined glycosylation sites from 40 controls and 70 patients in 14 inherited glycosylation-disorder groups after exclusion of four DYM-labelled samples without a prespecified target score. Group sizes ranged from 1 to 20 patients, with four single-patient groups (Fig 3A; Table S1).

We summarised each composition using five glycan features defined without fitting a reaction network. The Hex-rich/HexNAc2 feature indicated whether a glycoform contained two N-acetylhexosamine (HexNAc) residues and at least five hexose (Hex) residues. The branching-associated feature counted HexNAc residues beyond two. The antenna-associated Hex feature counted Hex residues beyond three, capped at the HexNAc excess beyond two, and was zero for glycoforms with no HexNAc excess. The sialylation and fucosylation features counted total sialic-acid residues (NeuAc plus NeuGc) and fucose residues, respectively. These features are composition proxies, not resolved branch counts or enzyme activities; the detailed definitions are in Methods, “Composition-derived pathway scores”, and S1 Text, Supplementary Methods S2.

For each feature, we multiplied its value for each glycoform by that glycoform’s within-site proportion and summed over glycoforms. For example, hypothetical proportions of 20%, 30% and 50% for glycoforms carrying zero, one and two sialic-acid residues would give a site sialylation score of 0 × 0.2 + 1 × 0.3 + 2 × 0.5 = 1.3. This is a calculation example, not an observed profile. We then averaged the site scores with equal weight over each participant’s eligible observed sites to obtain five participant-level composition scores; missing sites were omitted from this average.

To compare features measured on different scales, we subtracted each score’s mean among the 40 controls and divided by its control sample standard deviation, giving a *z* score. Positive values indicate higher scores than the control mean and negative values indicate lower scores. We averaged patient *z* scores within each disorder before ranking the five features (Fig 3B). Because related glycan-trait changes had already been reported in this cohort, the analysis tested retrospective concordance rather than independent phenotype discovery.

The target score and its expected direction for each disorder were fixed before group scores were inspected. Six disorders with direct biosynthetic defects (the mechanism-anchored groups) affected galactosylation (B4GALT1), branching (MGAT2 and SLC35A3), sialylation (NANS and SLC35A1) or fucosylation (SLC35C1) [20–25]. Six other disorders shared an established downstream hyposialylation phenotype (the phenotype-anchored groups), whereas MOGS and MAN1B1 were treated as boundary cases because composition-level annotation did not directly resolve their expected products [26–28]. These strata are summarised in Fig 3E; complete assignments and their biological basis are provided in Table S2.

Within each disorder, the five group-mean, control-standardised scores were ranked by absolute magnitude, and agreement with the prespecified direction was evaluated separately. A target score was counted as Top-1 or Top-2 concordant only when it changed in the expected direction and ranked first or within the top two, respectively. The target score was Top-1 concordant in 10 of 14 groups and Top-2 concordant in 13 of 14 (Fig 3C,E; Table S2). Among the six mechanism-anchored disorders, four were Top-1 concordant and all six were Top-2 concordant (Fig 3E). Top-1 and Top-2 are descriptive ranking summaries, not significance thresholds or validated diagnostic criteria. Each disorder contributed equally to these counts irrespective of sample size.

With patient data held fixed, we resampled the controls and recalculated standardisation and ranks. Top-2 concordance for the six mechanism-anchored disorders was retained in 83.9–100% of replicates, and analyses restricted to 29 commonly observed sites, adjusted for coverage or limited to NeuAc-based sialylation retained the overall pattern (Fig 3D,E; S1 Text, Fig S1B and Table S4). MOGS ranked second in the expected direction, whereas MAN1B1 ranked fourth and changed in the opposite direction, showing that the prespecified score did not capture the expected MAN1B1-associated change. Four of the six mechanism-anchored groups contained only one patient, and the six phenotype-anchored groups shared the same downstream proxy; the evidence therefore supports retrospective pathway concordance rather than population-level causal identification.

### Disease-associated shifts align with predicted loss-of-function directions

We assessed whether site-specific responses to reaction-rate changes were consistent with the direction and magnitude of disease-associated glycoform shifts.

#### Control-composition fitting and sensitivity calculation

To interpret site-specific glycoform shifts in terms of biosynthetic reactions, we first fitted each network to control compositions and calculated its local responses to rate changes (Fig 4A).

The primary site models grouped processing into five effective classes: mannose trimming (MAN), branching (MGAT), galactosylation (GALT), sialylation (SIAT) and fucosylation (FUT). At each site, we retained the glycoform classes represented by its reaction network (the graph-supported classes).

To establish a control reference for each site network, we used the evaluable profiles among the 40 controls. For each control, abundances of the graph-supported classes were normalised to sum to one; these profiles were then averaged with equal weight across controls to obtain the target composition 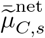 . Here, *C* denotes controls and *s* denotes the site. Controls with no observed profile or no graph-supported abundance at that site were omitted.

We held the network connections, entry state, secretion rates and observation classes fixed and adjusted the five effective rates, one for each reaction class, independently for each site. Each candidate rate vector *θ* specified reaction fluxes and hence a steady-state glycoform distribution. Using the model in “From reaction rates to secreted composition”, we aggregated the predicted secretion fluxes into the observed classes and normalised their sum to one to obtain the predicted proportions 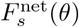.

We searched within the allowed rate ranges for a combination that minimised the sum of squared differences between these predicted proportions and the observed control mean:

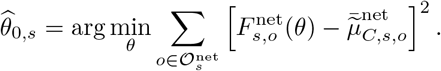

Here, 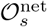 is the graph-supported class set and 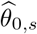 the fitted reference at site *s*. Patient profiles entered only the subsequent response comparison. This reference describes the control composition under the chosen network, without establishing unique rate recovery. Rate bounds and numerical optimisation are specified in Methods, “Control-composition fitting and model comparison”.

We then perturbed each reaction-class rate in turn around 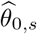, holding the other rates fixed, to calculate the local composition responses. These responses form the site-specific log-rate Jacobian *J*_*s*_, defined in “Model and definitions”. For small rate changes,

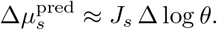

Column *J*_*s,r*_ gives the change in each glycoform proportion per unit increase in log rate *r*, holding the other rates fixed. Its positive and negative entries indicate which classes increase and decrease. Numerical sensitivity calculations are detailed in Methods, “Forward calculation and numerical sensitivities”.

#### Alignment with disorder-associated glycoform shifts

We first asked whether the observed disorder-associated shifts aligned with the composition responses expected from the affected reaction classes (Fig 4B,C).

For disorder contrast *d*, we defined the observed change and its directional agreement with the assigned reaction class *r* as

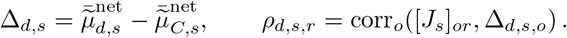

The first mean is over the observed patients in contrast *d*. The Pearson correlation is taken across glycoform classes *o* within the same site, comparing a sensitivity vector with the patient-minus-control change vector (Fig 4C). It is not a correlation across patients or between fitted and observed control abundances. Because the sensitivity represents a rate increase and the disorders reduce pathway function, agreement has a negative sign.

The observed redistributions aligned with the expected direction for all four reaction-class contrasts: across 30–35 evaluable sites, median correlations ranged from -0.63 to -0.15 under the primary rate bounds of 0.05–5 and the wider sensitivity bounds of 0.01–20 (Fig 4C; Fig S2; Table S5). Alignment was strongest for B4GALT1-associated galactosylation and weakest for MGAT2-associated branching.

For the non-network comparator, we replaced *J*_*s,r*_ in the within-site correlation with a glycan-feature vector calculated on the same glycoform classes, retaining the observed change Δ_*d,s*_. Using the same classes in both representations (a support-matched comparison) gave correlations of similar sign and magnitude, including -0.41 for B4GALT1 and -0.22 for MGAT2 (Fig 4C).

The MGAT2 contrast showed the weakest and least certain alignment: its protein-cluster resampling interval ranged from -0.390 to -0.022 across 30 sites (Fig 4C), and both MGAT2 and SLC35C1 were represented by a single patient. This interval therefore describes variation across measured proteins rather than patient-population uncertainty. These comparisons support an association between the observed shifts and the expected reactions.

We also asked whether sites with stronger model responses showed larger observed changes (Fig 4E). We computed the Pearson correlation between ∥*J*_*s,r*_∥_2_ and ∥Δ_*d,s*_∥_2_ across sites for each assigned contrast. These Euclidean norms measure response size, whereas the within-site correlations above compare response direction. The magnitude correlations changed when the allowed rate interval was widened from 0.05–5 to 0.01–20. Thus, directional alignment did not imply a stable association between network sensitivity and observed response magnitude across reference-rate bounds.

### Network responses support reaction-based reconstruction across sites

We next asked whether site-specific reaction-class responses could represent composition changes across sites. To assess reconstruction accuracy, we compared this network representation with a control-derived principal component analysis (PCA) basis of the same dimension.

To assess reconstruction across sites, we estimated a shared coefficient vector from all evaluable sites except the held site *s*, then used that site’s sensitivity matrix to reconstruct its change:

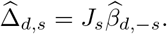

The predicted values 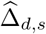 are changes in glycoform proportions. The vector 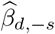 contains local alignment weights estimated from the other sites, not validated estimates of rate fold changes. For each disorder, Fig 4D reports the Pearson correlation between predicted and observed changes after concatenating the held-site class entries. The held site’s control fit and sensitivity matrix remain available, and the same patients contribute across sites, so this is conditional reconstruction rather than prediction in new patients. Details are in Methods, “Comparisons using the same observations and reconstruction across sites”, and S1 Text, Supplementary Methods S4, “Conditional reconstruction and PCA comparator”.

The network expressed cross-site composition changes as combinations of defined reaction-class responses. Its conditional leave-one-site-out reconstruction showed no consistent advantage over PCA with the same number of components (Fig 4D; Fig S3; Table S6).

### Most site models distinguish their rate parameters locally, but recovery depends on measurement precision

We next asked whether the measured glycoform classes could distinguish changes in individual reaction rates. If a small change in one rate produces the same composition response as a combination of changes in other rates, composition alone cannot separate those changes. We assessed this at the rates fitted to each site’s control composition, then examined how measurement error would limit the precision of recovery.

At 32 of 35 sites, the model responses distinguished small changes in all rate classes present in the network. To make this assessment, we retained the corresponding columns of the sensitivity matrix *J*_*s*_ and checked their independence by singular-value decomposition. These 32 matrices had full column rank, the local identifiability condition. The remaining three sites had five measured glycoform classes and five rate classes. Because five proportions summing to one supply only four independent values, these sites lacked enough information to distinguish all five rates. This assesses what the fixed model can distinguish near the fitted control rates; it does not demonstrate recovery of patients’ true rates. Numerical tolerances are given in Methods, “Local rank and conditioning”.

We then checked whether this result was specific to the fitted control rates. At each site, we generated 512 hypothetical rate settings by drawing each rate independently from a log-uniform distribution over 0.05–5 and evaluated the Jacobian rank at each setting. These were neither data-fitted rates nor optimisation starting points, and they need not reproduce the control composition. Of the 35 sites, 28 were full rank in every setting, four in 88.9–99.8% of settings, and the same three in none (Methods, “Local rank and conditioning”; S1 Text, Table S9). Thus, the rank result usually persisted beyond the fitted control point within the sampled settings; this finite check does not cover the whole continuous parameter range.

Even where rate changes were distinguishable, some combinations of changes produced much weaker composition responses than others. We summarised this difference by the condition number: the ratio of the strongest to the weakest response to log-rate changes of equal Euclidean length. Among the 32 full-rank sites, its median was 19.8, corresponding to an approximately 20-fold difference in response size. Seven sites had condition numbers above 100 and two above 1,000 (S1 Text, Table S8). These reference levels describe the spread of conditioning, not acceptable rate precision. Sites with more measured classes tended to have lower condition numbers (Spearman *ρ* = −0.36), but additional classes did not guarantee stable recovery.

We applied the noise-amplification calculation introduced in the canonical panel analysis (Fig 2) to the full-column-rank site models. Under the same measurement-error assumptions, we calculated each site’s factor *g*_*s*_ from its observed control mean composition and the sensitivity matrix at its fitted control rates, then used the same precision criterion to evaluate measurement requirements. The covariance calculation is detailed in Methods, “Measurement-error amplification”.

For the illustrative, analyst-selected target *δ* = 0.25, 24 sites would meet the target at an assumed CV of 5%, compared with 21 at 10% (S1 Text, Fig S4C and Table S13). These counts depend on the chosen target and CV; they are design scenarios, because the released data contained no technical replicates from which to estimate assay precision. Most site models therefore allowed rate changes to be distinguished locally, while the precision of their recovery depended on response strength and measurement error.

### Expanding from five to eight rate classes improves glycoform fit but reduces rate resolvability

Assigning separate rates to previously grouped reaction steps may improve composition fit, but also increases the number of rates to distinguish. We split mannose trimming into two stages (MAN1 and MAN2) and branching into three (MGAT1, MGAT2 and late-branch MGAT), while retaining GALT, SIAT and FUT, giving eight rate classes. The expanded model also allowed fucosylation independently of antenna addition, so the comparison included a change in reaction eligibility as well as rate-class separation (Methods, “Network construction”; S1 Text, Supplementary Methods S3).

We fitted each model separately to the control data at the same 35 sites, minimising the squared differences between predicted and mean observed proportions over each graph’s supported classes. The primary rate bounds were 0.05–5 for both models. We then compared the Pearson correlation between fitted and observed class proportions at each site. Its median increased from 0.221 to 0.938, and the number of sites with negative correlations fell from five to zero (Fig 5B). The gain persisted when the allowed rate interval was widened to 0.01–20 and across starting points, whereas graph coverage changed little (Methods, “Control-composition fitting and model comparison”; S1 Text, Table S10). Because both rate grouping and reaction eligibility changed, their separate contributions were not isolated.

We next evaluated the sensitivity matrix at each model’s own fitted control rates, using only the rate classes present in that site’s graph. Expanded sites contained five to eight such classes; full rank therefore required distinguishing the site’s actual rate set, not eight rates at every site. Despite the better composition fit, the number of full-rank sites decreased from 32 to 25 (Fig 5B). Among the full-rank sites in each model, the median condition number increased from 19.8 to 358.7. Equal-sized log-rate changes along the most and least sensitive directions thus differed about 359-fold in their composition response, compared with approximately 20-fold in the five-class model. The improved fit therefore did not imply more precise rate estimates.

We also repeated the conditional held-site reconstruction described above, estimating response coefficients from the other sites and reconstructing the excluded site’s composition change. PCA used five or eight components to match the corresponding network representation. Across disorders, the median reconstruction correlation changed from 0.394 to 0.471 for the network model and from 0.435 to 0.455 for PCA. Thus, reconstruction improved only modestly; these descriptive comparisons do not assess patient-level prediction (Methods, “Comparisons using the same observations and reconstruction across sites”; S1 Text, Table S10).

To examine why using weaker composition responses can increase estimation error, we conducted a separate simulation within the fixed canonical eight-rate model and a nine-class observation panel. We assumed a distribution of true log-rate changes and multinomial measurement noise. Singular-value decomposition ordered orthogonal combinations of log-rate changes (sensitivity modes) from strongest to weakest composition response. We varied how many modes were retained for estimation and calculated the expected squared error analytically: omitting modes introduced bias, whereas including weak modes amplified measurement-noise variance. Retaining more modes reduced squared bias but increased variance (Fig 5A). Under the illustrated noise setting, their sum was lowest at five retained modes; this optimum changed with measurement precision (S1 Text, Table S14). The simulation varied the estimated rate combinations within one model, rather than changing the biochemical reaction classes as in the empirical comparison. Simulation assumptions and error calculations are detailed in Methods, “Fixed-model truncation simulation”, and S1 Text, Supplementary Methods S4 and Supplementary Results S6a.

## Discussion

This study connects the biosynthetic information supported by site-specific glycoform compositions to the measurements needed to obtain it. Building on existing approaches to infer enzyme activities and assess parameter uncertainty [11–13], we evaluate how the choice of glycoform classes, measurement precision and model detail jointly determine which reaction-rate changes can be distinguished. Under a specified network and reference rates, this assessment helps researchers judge how far measured compositions support a biosynthetic explanation and design measurements to resolve additional reaction changes. Model fit remains essential, but must be considered alongside the ability to distinguish rates and the uncertainty introduced by measurement error.

A central finding is that improved composition fit can accompany reduced rate identifiability. Expanding the empirical model from five to eight reaction classes improved median fit correlation from 0.221 to 0.938, while reducing the number of full-rank sites from 32 to 25 and increasing noise sensitivity (Fig 5). More detailed models therefore require evaluation of both composition fit and rate uncertainty. The separate fixed-model simulation explains a related trade-off: retaining weak response directions reduces omitted-direction bias but amplifies noise. Its optimum depended on measurement precision, supporting selection of the detail to estimate according to the information available, without establishing a universally optimal number of reaction classes.

The plasma analysis distinguishes three levels of interpretation: detecting pathway-associated composition changes, relating them to reaction responses, and recovering individual rates. Expected composition scores showed Top-2 concordance in 13 of 14 disorder groups, and network responses aligned with the expected loss-of-function directions in the four reaction-class contrasts (Figs 3 and 4). Most site models also distinguished rate changes locally near their fitted control reference. This model property does not establish recovery of patients’ true rates. Direct composition features showed similar directional alignment, and reconstruction was comparable to PCA with the same number of components. The network provides a reaction-based interpretation and a way to assess distinguishability; these comparisons do not establish a predictive advantage specific to its topology.

The canonical analysis translates these distinctions into a measurement-design principle. At the specified reference rates, choosing the best qualifying nine-class panel allowed a 17.9-fold larger measurement CV than choosing the median panel for the same target rate uncertainty. Expanding the best nine-class panel to all 22 states reduced the best attainable noise amplification by less than 20% (Fig 2; Fig S4). Thus, panel composition can matter more than panel size under a specified model. Candidate classes can be compared by whether their responses separate the target reactions at an achievable measurement precision. Higher structural resolution may increase the number of observations but also the number of rates needed to represent them, so measurement resolution and model detail should be considered together.

Interpretation remains conditional on the model and reference rates. Primary-model fits were often poor, and all 35 fits reached at least one search bound (S1 Text, Table S8), limiting the biological interpretation of those references. Effective rates may combine tissue mixtures, substrate accessibility, donor supply, secretion and clearance. The well-mixed processing models omit structural, linkage and transport features, and the data lack relevant clinical covariates and technical replicates. These limitations affect model adequacy and uncertainty; full local rank alone cannot establish unique recovery over the full parameter range. The normalisation-based class-count bound remains a necessary condition for each independently fitted panel.

Further validation should test recovery of known rates on site-specific graphs under realistic measurement error, including cases where the fitted graph differs from the data-generating graph. Prospective technical replicates and clinical covariates would help separate analytical error from biological variation. The framework can guide these studies by identifying informative glycoform classes, setting measurement-precision targets and assessing how many reaction-rate changes the observations can support.

Together, these choices support biosynthetic interpretation at the level justified by the data and guide measurements to resolve additional reactions.

## Materials and methods

### Study data and analysis overview

We analysed the author-released, preprocessed PXD034214 plasma table of within-site glycoform percentages, rather than raw mass spectra or glycopeptide intensities.

Upstream spectrum processing is described by Wessels et al. [16]; the released table contained 68 peptide-defined sites, 40 controls and 74 candidate patients. We excluded four DYM-labelled samples without a prespecified target score, leaving 70 patients in 14 disorder groups. Group sizes ranged from 1 to 20, with four single-patient groups (S1 Text, Tables S1 and S2). For sample *i*, peptide-defined site *s* and released glycoform row *r*, percentages *x*_*i,s,r*_ were expressed as within-site proportions,

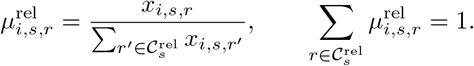

Here, 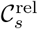 is the set of glycoform rows reported for site *s*. This operation only changes the scale from percentages to proportions. Profiles with zero site sum were treated as missing and were not imputed. We used two complementary representations. Composition-derived pathway scores summarised predefined glycan features using the complete released compositions. A reaction-network model instead mapped effective biosynthetic rates to graph-supported compositions and quantified local response directions around a fitted control reference.

Each released glycan composition was written as *g* = (*H, N, F, A, G*), containing the counts of hexose (Hex), N-acetylhexosamine (HexNAc), deoxyhexose (dHex),N-acetylneuraminic acid (NeuAc, also written Neu5Ac) and N-glycolylneuraminic acid (NeuGc). In this plasma N-glycan analysis, dHex was treated operationally as fucose; composition annotation does not distinguish all deoxyhexoses.

The data were first accessed approximately in April 2026; the exact initial access date was not recorded. The deposited filename and checksum were verified against the public PRIDE record on 5 October 2026. This study was a secondary analysis of public,de-identified data. We recruited no participants, collected no specimens and accessed no direct identifiers. Ethical approval and consent procedures for the original study are reported by Wessels et al. [16].

### Composition-derived pathway scores

#### Score calculation and concordance

To assess pathway-associated composition changes without fitting a reaction network, we defined five glycan features associated with the pathways of interest,

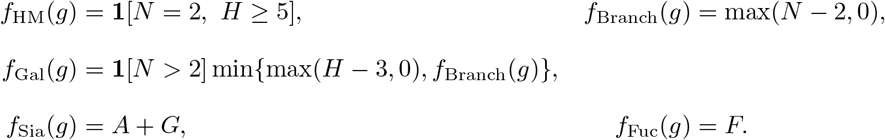

Here, **1**[·] equals one when the stated condition holds and zero otherwise.

For each feature *e*, the composition-weighted mean was calculated at each observed site and then averaged equally across the eligible sites *S*_*i*_ in sample *i*,

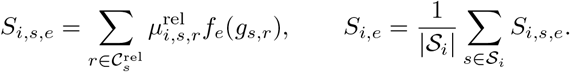

Here, *e* indexes the five features above. The resulting scores are pathway proxies, not structure-resolved counts or estimates of individual enzyme activity. In particular, HexNAc and Hex do not uniquely identify branching and galactose in all glycan classes; linkage, branch position and core versus antenna fucosylation were unresolved. NeuAc and NeuGc were summed computationally without implying biochemical equivalence. Eligible sites were those with a positive released-profile sum in that sample; missing sites contributed neither a zero score nor a denominator term. Composition scores used all such released sites independently of network inclusion. Additional annotation details are provided in S1 Text, Supplementary Methods S1–S2.

Each patient’s score was standardised to the mean and sample standard deviation of the 40 controls. Within each disorder, the five absolute group-mean standardised scores were ranked. A prespecified target score was called Top-1 or Top-2 concordant only when it had the expected sign and ranked first or within the first two, respectively. Disorders, rather than patients, contributed equally to aggregate counts. The target score and direction for each disorder were fixed before group scores were inspected. The subsequent organisation into mechanism-anchored, phenotype-anchored and boundary-case strata was performed after inspection, so stratum-specific summaries are post hoc and descriptive. The study was retrospective and was not preregistered.Assignments and group sizes are in S1 Text, Table S2; the supplementary assignment-enumeration reference is described in Supplementary Methods S5.

#### Control calibration and coverage sensitivity

For control-calibration uncertainty, we generated 1,000 bootstrap samples of 40 controls with replacement (seed 20260713), using the same resamples for all disorders. Patient scores, target assignments and expected directions were held fixed. Each resample supplied new control means and sample standard deviations, from which we recalculated standardised group means, absolute-score ranks and directional Top-1/Top-2 concordance. The reported frequencies quantify stability of control calibration, not patient-population uncertainty.

We separately repeated the score analysis using only the 29 sites observed in all 40 controls and 70 patients. Scores were recomputed over this common set and restandardised before ranking. In a coverage adjustment, each original participant-level score was regressed on detected-site count with an intercept using controls only. The fitted control relationship was subtracted from every participant’s score; residuals were standardised to the control residual mean and sample standard deviation before repeating the same ranking and direction checks. A NeuAc-only sensitivity replaced *A* + *G* by *A* in the sialylation score. These analyses did not impute missing sites (S1 Text, Supplementary Methods S2 and S5; Supplementary Results S7 and Table S4).

### Reaction networks and the forward model

#### Network construction

For each site *s*, we generated a directed graph of reachable glycan states using the five primary reaction classes defined in Results. The latent state tracked excess mannose above the Man3 core, added HexNAc antennae, galactose and sialic-acid counts, and binary fucose presence. State ranges were determined from the site’s observed compositions and entry-state requirements. Before the first antenna was added, mannose trimming stopped at Man5; the first MGAT step was allowed at Man5, and subsequent branching required trimming to the Man3 core. Galactosylation and sialylation were limited by the numbers of available antennae and galactose residues, respectively. The primary model allowed fucosylation only after the first antenna had been added. States unreachable from the input were removed, and the remaining states were mapped to observed compositions.

For the computational input, we identified classes with two HexNAc residues and at least five Hex residues, selected their largest Hex count, and injected unit input into the corresponding unprocessed mannose-rich state. If no such class was present, the input was Man5GlcNAc2. This rule truncates the computational network to the observed support; it does not infer the biological glycan-transfer substrate. Complete state definitions and reaction-specific eligibility tests are given in S1 Text, Supplementary Methods S3 and the supplied code.

To assess the effect of adding reaction classes, an eight-class model separated MAN into MAN1 and MAN2 and MGAT into MGAT1, MGAT2 and a late-branch MGAT (MGAT4/5), while retaining lumped GALT, SIAT and FUT classes. Unlike the primary graph, it allowed composition-level FUT reactions independently of antenna addition. This rule can represent fucose-positive HexNAc2 compositions but does not specify a fucose position or a structure-specific FUT8 substrate (S1 Text, Supplementary Methods S3).

#### Forward calculation and numerical sensitivities

We used the reaction matrix and steady-state equations defined in “From reaction rates to secreted composition”, with site-specific matrices and vectors indexed by *s*. The canonical calculation solved the linear system directly, with unit input at Man9 and uniform secretion *κ*_*i*_ = 0.5. For the empirical acyclic graphs, states were ordered so that every reaction pointed from an earlier to a later state. With *κ*_*i*_ = 1, forward substitution then solved the same balance equations, passing each state’s reaction fluxes to its downstream states. This avoids repeated dense solves. The system has a unique non-negative solution for admissible rates; the proof is given in S1 Text, Supplementary Methods S3.

The site-specific aggregation matrix *B*_*s*_ retained graph-supported observed classes. Their normalised secretion fluxes defined 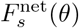, the site-specific counterpart of the panel composition *µ*_*P*_ (*θ*). The supported observation set was held fixed during rate perturbations. Mapping and normalisation of empirical observations are detailed in “Observed classes and conditional compositions” and S1 Text, Supplementary Methods S3.

For canonical panel enumeration, we approximated log-rate derivatives by forward differences with step 10^*−*6^, then differentiated the within-panel normalisation. For empirical site models, each rate was multiplied by 1 + *ϵ* and 1 − *ϵ*, with *ϵ* = 10^*−*3^; the two renormalised compositions were differenced and divided by 2*ϵ* to approximate the log-rate derivative. These calculations include the effect of renormalisation rather than differentiating unnormalised flux alone. Rank thresholds are specified under “Local rank and conditioning”; canonical stability checks are in S1 Text, Supplementary Results S6, and empirical checks are in Tables S8 and S9.

### Observation mapping and model fitting

To compare measured and predicted compositions, we first mapped the released glycoforms to the observation classes represented by each network. We then fitted the rates using the mean control composition on that shared set of classes.

#### Observed classes and conditional compositions

Released compositions were coarsened as

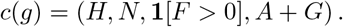

The primary mapping required *H* ≥ 3 and *N* ≥ 2, and excluded fucose-positive HexNAc2 rows that the primary reaction rules could not represent. We required at least five distinct candidate classes before graph construction and at least five reachable observed classes after pruning, together with an evaluable control mean. A computational size guard excluded graphs with more than 600 reachable states. Of the 68 released sites, 33 had only one to four candidate classes; the remaining 35 met all criteria and retained 5–36 observed classes, with at most 277 latent states. The five-class inclusion cutoff was an analysis setting, not the *m* ≥ *k* + 1 condition for identifying all active rates. The augmented model used these 35 sites with its own supported classes.

For observed class *o*, the proportions of its constituent released rows were summed to *z*_*i,s,o*_, restricted to the graph-reachable set 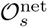, and conditionally normalised,

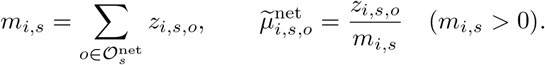

Thus the network target was the graph-supported conditional composition, not the complete released composition. The excluded abundance fraction, 1 − *m*_*i,s*_, was recorded as unsupported mass. Profiles with *m*_*i,s*_ = 0 contained no abundance represented by the network and could not be evaluated. Complete latent-state definitions, reaction rules, entry-state choices and coverage diagnostics are given in S1 Text, Supplementary Methods S3.

#### Control-composition fitting and model comparison

For each site, we first normalised each evaluable control profile over its graph-supported classes and then averaged these profiles with equal weight. Controls with no observed profile or no graph-supported abundance at that site were omitted. The resulting mean, 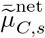, was fitted independently at each site:

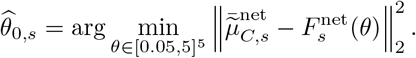

Here, *C* denotes controls. At each candidate rate vector, the forward calculation produced steady-state secretion fluxes, which were aggregated and normalised over the same observed classes as the target. The objective was an unweighted sum of squared class-proportion residuals, with no rate penalty. Patient compositions were not used to fit this reference. The primary and expanded rate bounds are computational search ranges, not measured physiological limits; fitted rates and downstream sensitivities remain conditional on them.

We optimised *η* = log *θ* with the bounded limited-memory Broyden–Fletcher–Goldfarb–Shanno algorithm (L-BFGS-B) in SciPy. For the primary five-class model, six starting vectors were drawn from independent standard normal log-rate distributions using a fixed random-number sequence (seed 0), then clipped to [log(0.05), log(5)]. Each run allowed at most 400 iterations; other solver settings used the defaults of the software version recorded in S1 Text, Table S7. We retained the returned vector with the smallest squared-error objective across the six runs. This procedure selects the best solution found, without establishing a unique or global optimum. The primary implementation optimised all five coordinates; any graph-absent rate had no effect on the objective and was excluded from active-column identifiability diagnostics. The retained vector defined the operating point for subsequent sensitivities, not a directly measured biochemical state. Fit was assessed by squared error and the Pearson correlation between predicted and mean observed control proportions across classes.

The augmented eight-class model was fitted to the same observations at the same 35 sites. Only graph-present rates were optimised; absent classes were fixed at one and excluded from boundary counts, rank denominators and condition numbers. Its 36 starting points comprised one unit-rate vector and 35 standard normal log-rate vectors (seed 20260713), clipped to the same bounds. L-BFGS-B allowed 800 iterations, with relative objective tolerance 10^*−*12^ and projected-gradient tolerance 10^*−*8^; the lowest-objective returned vector was retained. Increasing from 12 to 36 starts changed no site’s objective by more than 10^*−*8^. For both models, the complete fitting and downstream analysis was repeated with bounds [0.01, 20] per rate. We compared fit, active-column rank, conditioning and conditional reconstruction; the comparison results are in S1 Text, Table S10.

### Local identifiability and measurement precision

#### Local rank and conditioning

The class-count requirement applies separately to each jointly normalised panel and depends on the rate classes present in its network. Information from other sites does not alter this requirement when sites are fitted independently. A model with parameters shared across sites would require a rank analysis of the combined observations.

We computed the Jacobian rank and condition number from the singular values of the log-rate Jacobian. For a full-column-rank Jacobian, the condition number is the largest singular value divided by the smallest.

The canonical graph contained 22 states and 25 reactions grouped into eight rate classes. We evaluated its full output and exhaustively enumerated renormalised state subsets. Panels were called full rank when the eighth singular value exceeded 10^*−*9^, with canonical tolerance and operating-point checks reported in S1 Text, Supplementary Results S6 and Table S3. This is a numerical tolerance for treating small singular values as zero, not a biological precision requirement. The dimensional bound *m* ≥ *k* + 1 follows from composition normalisation and does not depend on this tolerance. Empirical active-column rank used the same absolute threshold (Table S8).

To assess dependence on the fitted reference, we drew 512 five-component rate vectors with independent log-uniform components over [0.05, 5], and another 512 over [10^*−*3^, 10^3^] (seed 20260713). The same vectors were evaluated at every site, retaining only graph-present reaction classes for rank calculations. We recomputed the Jacobian without changing the graph or refitting to observations. These settings were hypothetical evaluation points, not optimiser starts or fitted rates. The wider range was a numerical stress test; the sampling count and distributions were analysis choices. Fractions of full-rank settings describe the sampled points and do not establish rank over the continuous range (S1 Text, Table S9).

#### Measurement-error amplification

For a panel composition *µ*_*P*_, independent relative measurement error with coefficient of variation CV gives the first-order composition covariance

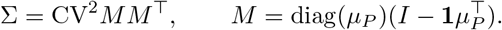

Here, *I* is the identity matrix and **1** is a vector of ones. Canonical panels used the model-predicted composition; empirical panels used the observed graph-supported control mean and the Jacobian at the fitted control reference, retaining only graph-present rate classes. For full-column-rank *J*_*P*_, where the resulting information matrix is invertible, the covariance of generalised least-squares estimates in the local linear model is 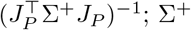 denotes the Moore–Penrose pseudoinverse. We set CV = 1 and took the square root of this covariance matrix’s largest eigenvalue to define *g*, the maximum standard deviation over unit-length log-rate directions. For target *δ*, CV ≤ *δ/g*.

Numerically, we used singular-value decomposition to scale sensitivity directions by their measurement uncertainty and invert only retained directions. We assigned *g* = ∞ when the rank or inversion criteria failed. Thirty empirical sites had finite *g*; two of the 32 full-rank sites had too few positive-mean abundance directions under these criteria. Therefore, full Jacobian rank alone did not guarantee a finite precision requirement under the assumed relative-error model. The SVD-based numerical evaluation and its thresholds are specified in S1 Text, Supplementary Methods S4, “Numerical evaluation of measurement-error amplification”. Tables S11 and S12 report canonical precision calculations; Table S13 reports empirical requirements and the nonlinear recovery check.

### Reaction attribution and conditional reconstruction

#### Local-response attribution

At each site *s*, we applied the local-response definition in Model and definitions to 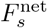, evaluating its log-rate Jacobian *J*_*s*_ at the control fit 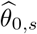.

For disorder *d*, evaluable patient profiles were normalised separately and averaged with equal patient weight. The observed site response was the difference between this patient mean and the observed control mean, rather than the fitted control prediction.

We denoted this patient-minus-control difference by 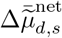, abbreviated as Δ _*d,s*_ in Results. We assessed whether the response aligned oppositely with the positive-rate sensitivity expected for a loss-of-function contrast. We stacked the observed responses and Jacobians across sites into 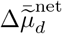 and *J*, then estimated a coefficient vector by ridge regression,

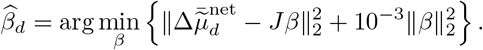

The entries of this unconstrained coefficient vector are local alignment weights, not estimated fold changes or absolute rates. We applied no additional site weights. The SIAT contrast pooled the six NANS and one SLC35A1 patients with equal patient weights; equal disorder weights were examined separately. Disorder–site pairs with missing data or no abundance represented by the network were omitted rather than replaced by zero.

#### Comparisons using the same observations and reconstruction across sites

Within each site, we calculated the Pearson correlation across graph-supported classes between the assigned network sensitivity and the observed loss-of-function response. Both vectors were centred over classes; pairs with zero variance in either vector were excluded. For the non-network comparator, a coarsened class (*H, N, b, S*) had feature values max(*N* − 2, 0) for MGAT, min{max(*H* − 3, 0), max(*N* − 2, 0)} for GALT, *S* for SIAT and *b* for FUT, where *b* = **1**[*F >* 0] and *S* = *A* + *G*. Each feature was centred over the same classes and correlated with the same observed response. Thus FUT represented binary fucose presence here, whereas the pathway score counted fucose residues. For the magnitude comparison, we correlated the Euclidean norms of sensitivity and observed response across sites.

Descriptive 95% intervals used 2,000 bootstrap resamples of the 21–24 measured proteins, sampling proteins with replacement and retaining all their sites together; interval limits were the 2.5th and 97.5th percentiles. For sign-flip tail fractions, each protein’s site correlations received a shared random sign in each of 10,000 draws. We counted draws whose median was no greater than the observed median and added one to both the count and denominator. This reference assumes joint sign symmetry within proteins. These summaries were not multiplicity-adjusted and do not estimate patient- or proteome-population uncertainty.

For conditional leave-one-site-out reconstruction, we retained the held site’s control fit and basis and estimated the disorder-specific coefficient from the other evaluable sites:

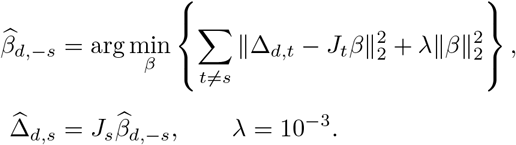

Here, *s* is the held site, *t* indexes the training sites, and the ridge penalty stabilises the coefficient estimate. We fixed *λ* = 10^*−*3^ as an analysis setting shared by the network and PCA comparisons; it was not selected by predictive validation, and reconstruction results are conditional on this choice. The held site’s disorder response was excluded from this fit. We centred each basis column over the site’s classes, stacked training-site bases into *X* and responses into *y*, and solved 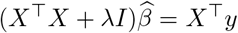. For each disorder, held-site predictions and observations were concatenated over all evaluable sites and classes before calculating their Pearson correlation. Consequently, sites with more classes contributed more entries; multi-disorder summaries gave equal weight to disorder-specific correlations.

For the principal component analysis (PCA) comparator, we formed the union of graph-supported observed class keys. Each control’s conditional site profiles were mapped to these keys, summed and divided by that control’s number of observed modelled sites. Missing sites were omitted from both the sum and denominator. We centred the resulting 40 control vectors by subtracting their column means and retained the first five right singular vectors, or eight for the augmented comparison. At each site, the corresponding class rows defined its PCA basis; columns were centred within that site and entered the same ridge reconstruction with the same penalty and observations. This matches basis dimension, but PCA supplies a global basis whereas the network supplies separately fitted site-specific bases.

The same patients contributed responses at training and held sites; this analysis therefore evaluates within-disorder reconstruction across sites, not prediction in new patients or cohorts. Additional comparator limitations are discussed in S1 Text, Supplementary Methods S4, “Conditional reconstruction and PCA comparator”. The 90% and 95% graph-supported-mass filters are analyst-selected coverage sensitivity checks, not validated adequacy thresholds. Graph-coverage filters and rate-bound sensitivity results are reported in S1 Text, Supplementary Results S8 and Tables S5 and S6.

### Fixed-model truncation simulation

We separately simulated the bias–variance trade-off of regularised inversion within the fixed canonical eight-rate model. Starting from an empty panel at unit reaction rates, a deterministic greedy procedure added the state that maximised the current panel rank, breaking ties in the fixed state order, until rank eight was reached. The resulting nine-state panel was held fixed. We drew 2,048 reference vectors log *θ* ∼ *N* (0, 0.7^2^*I*_8_) (seed 20260827); at each vector, we computed the panel composition and its Jacobian using central log-rate differences of 10^*−*5^.

Truncated singular-value estimators retained *r* = 1, …, 8 orthogonal log-rate directions (sensitivity modes), ordered by the strength of their composition responses. True log-rate changes were assumed to have zero mean and isotropic covariance 0.25^2^*I*_8_. To represent the noise in a difference between two composition measurements, we used two independent multinomial measurements, each with total count *N*_eff_ . Their difference has covariance 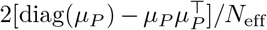under the local reference composition. We propagated this covariance through each truncated inverse and added the squared bias from omitted directions to obtain expected squared rate error, normalised by the expected squared true log-rate change. We summarised medians across reference vectors. No noisy measurement replicates or patient rate estimates were generated.

The effective count is the assumed total count in the multinomial model and determines the composition noise. The simulation shown in Results used 300,000; counts from 10^3^ to 10^6^ were examined in Table S14. The operating-point spread of 0.7 and effect standard deviation of 0.25 are also analyst-selected simulation assumptions, not estimates from patients; the latter is distinct from the target error *δ*. These are simulation design values, not empirical read counts, and the retained-mode count is distinct from the empirical model’s number of reaction classes. The selected panel and full simulation equations are in S1 Text, Supplementary Methods S4, “Idealised truncated-SVD bias–variance simulation”; the corresponding results are in S1 Text, Supplementary Results S6a and Table S14.

### Numerical implementation and reproducibility

All calculations were implemented in R and Python. Software versions, optimisation settings, tolerances and seeds are recorded in S1 Text, Table S7; the input identity and checksum are given in Supplementary Methods S1. The reproducibility archive on Zenodo (v1.1.0; https://doi.org/10.5281/zenodo.23176011) contains the complete analysis code, reaction rules, mappings, fixed parameters and machine-readable outputs. Original code is released under the MIT License.

A companion R package, `glycoPathDesign`, provides an interactive Shiny interface for exploring specified reaction networks, mappings from network states to measured classes, measurement-panel design and effective-rate fitting. The pathway display distinguishes selected rates from fixed reference rates and reports pooled measurements as class totals. Glycan drawings and enzyme-style identifiers encode model assumptions; they do not establish structures or enzyme activities from compositional data. Fit convergence, local rank and precision are assessed separately. This exploratory implementation is distinct from the R/Python pipeline used to generate the results reported here, and its default fitting settings do not define the study’s fitting protocol. The companion package is publicly available as release v0.6.0 at https://github.com/matsui-lab/glycoPathDesign; the hosted application is accessible without login at https://igcore.cloud/glycoPathDesign/. Its source and the standalone deployment source are included in the Zenodo archive cited above.

### Use of artificial-intelligence-assisted tools

OpenAI Codex assisted code development and auditing, figure preparation, manuscript drafting and revision, and Japanese translation. OpenAI image generation was used for non-data conceptual artwork and workflow illustrations in Figs 1 and 3. Checks included comparison of reported values with archived numerical outputs, an end-to-end rerun of the supplied numerical pipeline, verification of references against source records, and inspection of figure and document layouts. Experimental data were obtained from the public source study; AI-generated illustrations do not represent experimental observations. Available artwork source records, assets and file hashes are included in the Zenodo reproducibility archive. The authors are responsible for the scientific content and interpretation.

## Data availability

The experimental data analysed in this study are publicly available without restriction from PRIDE under accession PXD034214 (https://www.ebi.ac.uk/pride/archive/projects/PXD034214), released by Wessels et al. [16]. No special access privileges were required. The source workbook is distributed by PRIDE; its identity, checksum and provenance are documented in S1 Text and the reproducibility archive on Zenodo (v1.1.0; https://doi.org/10.5281/zenodo.23176011). This archive contains the complete analysis code, environment specification, run manifest, machine-readable results, figure assets and verification records, including the observation-panel design, P15 disease-label sensitivity, per-site identifiability, augmented-model and bias–variance analyses. The companion application is available at https://igcore.cloud/glycoPathDesign/, with source code at https://github.com/matsui-lab/glycoPathDesign (v0.6.0). Original code is released under the MIT License.

## Supporting information

**S1 Text. Supplementary Information**. Data provenance, reaction rules, numerical methods, fixed mappings, exact assignment enumeration, coverage sensitivity, within-site alignment, conditional reconstruction and reproducibility diagnostics, with Tables S1-S14 and Figures S1-S4.

## Acknowledgments

We thank the depositors of the public glycoproteomic datasets analysed in this study.

